# ThermoFinder: A sequence-based thermophilic proteins prediction framework

**DOI:** 10.1101/2024.01.02.573852

**Authors:** Han Yu, Xiaozhou Luo

## Abstract

**Motivation:** Thermophilic proteins are important for academic research and industrial processes, and various computational methods have been developed to identify and screen them. However, their performance has been limited due to the lack of high-quality labeled data and efficient models for representing protein. Here, we proposed a novel sequence-based thermophilic proteins prediction framework, called ThermoFinder.

**Results:** In this study, we demonstrated that ThermoFinder outperforms previous state-of-the-art tools on two benchmark datasets, and feature ablation experiments confirmed the effectiveness of our approach. Additionally, ThermoFinder exhibited exceptional performance and consistency across two newly constructed datasets, one of these was specifically constructed for the regression-based prediction of temperature optimum values directly derived from protein sequences. The feature importance analysis, using shapley additive explanations, further validated the advantages of ThermoFinder. We believe that ThermoFinder will be a valuable and comprehensive framework for predicting thermophilic proteins.

## 1. Introduction

Thermophilic proteins, also known as heat-stable proteins, are a class of proteins capable of retaining their structure and functionality under high-temperature conditions [1]. These proteins are typically obtained from thermophilic bacteria, particularly archaea, and have gained considerable significance in various fields such as biotechnology, food processing, and pharmaceuticals, owing to their enhanced protein stability, augmented enzymatic activity, and feasibility for high-temperature industrial processes [2–3]. For instance, thermally stable cellulases from thermophilic bacteria have been employed for the efficient conversion of lignocellulosic biomass to biofuels [4]. However, the identification and screening of thermophilic proteins is a demanding and laborious process [5]. To accelerate the progress of related fields, it is crucial to develop a high-throughput screening method for the rapid prediction of thermophilic proteins.

Computer-aided methods have made significant progress in identifying protein or peptide function owing to their speed, accuracy, and scalability advantages [6–9]. These methods have been widely employed in predicting thermophilic proteins, which are essential for discovering new thermophilic proteins and designing protein engineering strategies. Changli *et al* developed a novel method that combines mixed features and machine learning to recognize thermophilic proteins, while Zifan *et al* utilized feature dimension reduction technology and a Library for Support Vector Machines (LIBSVM) for the identification of thermophilic proteins [10–11]. Moreover, Phasit *et al* introduced a new computational method, SAPPHIRE, for more precise identification of thermophilic proteins using only sequence information without requiring structural information [12]. Additionally, Chaolu *et al* constructed an Support Vector Machine (SVM)-based identifier called TMPpred to differentiate between thermophilic and non-thermophilic proteins [13]. More recently, Jianjun *et al* proposed a deep learning model based on self-attention and multiple-channel feature fusion to predict thermophilic proteins, called DeepTP [14]. These methods primarily rely on intricate hand-crafted features. Nonetheless, accurately predicting thermophilic proteins still presents numerous challenges, such as the inadequacy of high-quality labeled data and the complex relationships between protein sequences and their functions.

To address these issues, we need to leverage more data and more sophisticated models to bridge the gap between protein sequences and their functions. Deep learning techniques, particularly self-supervised learning models, have recently made significant progress in natural language processing [15–16]. Self-supervised learning is a type of machine learning in which a model is trained to make predictions about a dataset without any explicit human-labeled supervision [17]. Similarly, this approach has been utilized to learn protein representations from unsupervised sequences. Numerous self-supervised methods have been employed to generate protein representations and enhance downstream task performance [18]. For example, Maxwell *et al* trained deep learning models based on convolutional neural networks to precisely predict functional annotations for unaligned amino acid sequences across rigorous benchmark assessments built from the 17,929 families of the protein families database Pfam [19]. Ahmed *et al* trained two auto-regressive models (Transformer-XL, XLNet) and four auto-encoder models (BERT, Albert, Electra, T5) on data from UniRef and BFD containing up to 393 billion amino acids, significantly improving per-residue and per-protein task predictions [20]. Felix *et al* utilized protein language models and conditional random fields to accurately predict various signal peptides, resulting in state-of-the-art results [21]. Moreover, the combinations of representations generated by self-supervised methods have been demonstrated to be effective in predicting protein structure and function [22–23]. However, there is still much to explore in terms of combining embedded representations generated by different types of self-supervised learning networks and comprehensively validating their performance.

Here, we developed ThermoFinder, a novel sequence-based framework for predicting thermophilic proteins. Our approach outperformed existing tools on two benchmark datasets, and we validated its effectiveness through ablation experiments. To further confirm the advanced performance of ThermoFinder and its unique ability for regression tasks, we constructed two new datasets, ThermoSeq_c 1.0 for classification and ThermoSeq_r 1.0 for regression, with a significantly larger number of experimentally validated data points. Furthermore, we analyzed the feature importance of four datasets using the SHapley Additive exPlanations algorithm (SHAP). Our findings highlighted the potential of ThermoFinder to improve the identification of thermophilic proteins from sequence data.

## 2. Materials and methods

### 2.1 Benchmark datasets

In order to validate the effectiveness of the proposed model, we selected two benchmark datasets. The first dataset, constructed by Zahoor *et al*, contained 1,364 thermophilic and 1,440 non-thermophilic proteins [24]. The second dataset, a collection combining previously reported datasets, contained 1,853 thermophilic and 1,853 non-thermophilic proteins [25]. The division of these two datasets follows the guidelines described in these two references and the resulted training and test sets are kept as the same for all training and test processes [24–25].

As the size of these datasets is relatively small, and they are both binary classification problems, we constructed two additional datasets, ThermoSeq_c 1.0 and ThermoSeq_r 1.0, for more reliable and comprehensive validation. ThermoSeq_c 1.0 was created as follows:

i. We obtained the optimal growth temperature of 21,498 microorganisms from Martin’s established dataset and selected the top 79 bacterial organisms with optimal growth temperatures greater than or equal to 70 ℃ as positive samples [26]. Similarly, we selected 79 organisms with optimal growth temperatures less than or equal to 30 ℃ as negative samples.
ii. For both the positive and negative group, we identified and downloaded 58 available proteomes from Uniprot database based on organism name, and obtained protein sequences from these proteomes [27].
iii. To remove redundancy and homology bias, and ensure that there are no identical or highly similar proteins in both the training and test sets, we combined all sequences from the two proteome categories and used the CD-HIT program to cluster similar sequences with a cutoff sequence identity of 0.4, resulting in 31,962 positive and 115,642 negative samples [28].
iv. To ensure an unbiased model, we randomly selected 31,692 negative samples to match the number of positive samples. All samples were randomly divided into 80% training and 20% testing sets.

Additionally, we also established a regression dataset, ThermoSeq_r 1.0, using the following procedure:

i. We obtained the temperature optimum of 21,774 proteins and their primary accession numbers from the BRENDA database, removing sequences without accurate temperature optimum or primary accession numbers [29].
ii. Then we downloaded all 7,348 available protein sequences from Uniprot database based on the primary accession numbers [27].
iii. We combined these sequences with their corresponding labels, temperature optimum, and randomly divided all samples into 80% training and 20% testing sets.

Here, we have removed sequences containing non-natural amino acids, such as ’XBUJZO’. Additionally, proteins with excessively long sequences exceeding 1000 were truncated to 1000 as instructed in previous publications [30]. The information of these datasets is listed in Table 1.

**Table1.**
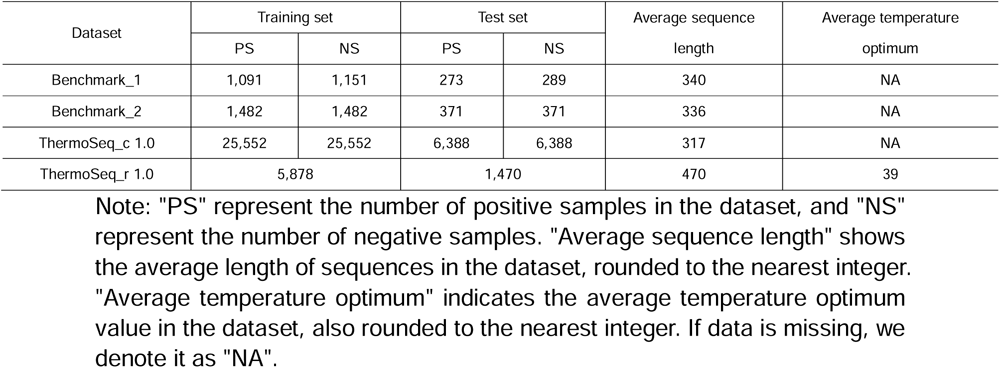
Information on four thermophilic proteins datasets.

| Dataset | Training set |  | Test set |  | Average sequence length | Average temperature optimum |
| --- | --- | --- | --- | --- | --- | --- |
|  | PS | NS | PS | NS |  |  |
| Benchmark_1 | 1,091 | 1,151 | 273 | 289 | 340 | NA |
| Benchmark_2 | 1,482 | 1,482 | 371 | 371 | 336 | NA |
| ThermoSeq_c 1.0 | 25,552 | 25,552 | 6,388 | 6,388 | 317 | NA |
| ThermoSeq_r 1.0 | 5,878 |  | 1,470 |  | 470 | 39 |

Note: "PS" represent the number of positive samples in the dataset, and "NS" represent the number of negative samples. "Average sequence length" shows the average length of sequences in the dataset, rounded to the nearest integer. "Average temperature optimum" indicates the average temperature optimum value in the dataset, also rounded to the nearest integer. If data is missing, we denote it as "NA".

### 2.2 Protein sequence representation

To represent protein sequences, we utilized four representative approaches: SeqVec, ProtCNN, ProtTrans, and CPCProt, as indicated in Table 2. These approaches generate embedded vectors that leverage unsupervised information to create effective protein representations that are easy for machine learning models to learn from. Below, we describe each approach in more detail.

**Table2.** Information on four protein representations.

| Features | Algorithm | Dimension |
| --- | --- | --- |
| SeqVec | LSTM (ELMO) | 1,024 |
| ProtCNN | CNN | 1,100 |
| ProtTrans | Transformer | 1,024 |
| CPCProt | Contrastive predictive coding | 1,536 |

#### 2.2.1 SeqVec representation

SeqVec utilizes the language model ELMo (Embeddings from Language Model) sourced from natural language processing to generate continuous vectors that represent protein sequences [31]. ELMo is a dynamic language model implementation that has been effective in per-residue and per-protein tasks. The resulting embedded vectors are 1,024 dimensions. SeqVec is available in the data repository https://github.com/rostlab/SeqVec.

#### 2.2.2 ProtCNN representation

ProtCNN is a deep learning model that uses convolutional ResNets to predict which Pfam family a given protein sequence belongs to [19]. This approach is based on convolutional neural networks that are commonly used in computer vision tasks. ProtCNN uses dilated convolution on protein sequences trained with the whole Pfam database to generate embedded vectors of 1,100 dimensions, which can be accessed in the data repository https://github.com/google-research/google-research/tree/master/ using_dl_to_annotate_protein_universe.

#### 2.2.3 ProtTrans representation

ProtTrans utilizes a series of transformer-based models, including auto-regressive models (Transformer-XL, XLNet) and auto-encoder models (BERT, Albert, Electra, T5), on data from UniRef and BFD containing up to 393 billion amino acids [20]. These models utilize a self-attention mechanism that allows them to weigh the importance of different parts of the input sequence and focus on the most relevant information. ProtTrans has achieved state-of-the-art (SOTA) results on several per-residue and per-protein prediction tasks. In this case, we utilized ProtT5-XL-UniRef50 to generate a 1,024-dimensional embedded vector. ProtTrans is available in the data repository https://github.com/agemagician/ProtTrans.

#### 2.2.4 CPCProt representation

CPCProt is a model that maximizes mutual information between context and local embeddings by minimizing a contrastive loss [32]. This approach is based on contrastive learning and aims to learn useful representations of data by contrasting similar and dissimilar examples. CPCProt has achieved comparable performance to SOTA self-supervised models for protein sequence embeddings on various downstream tasks while reducing the number of parameters. This model is the first to use contrastive predictive coding for protein representation and generates embedded vectors of 1,536 dimensions, which can be accessed in the data repository https://github.com/amyxlu/CPCProt.

### 2.3 Proposed framework

A machine learning framework was developed for predicting thermophilic proteins based on protein sequences. The framework consists of four steps: a dataset construction step, a feature engineering step, a model training step and a model evaluation step. This framework is shown in Figure 1.

**Figure 1.**
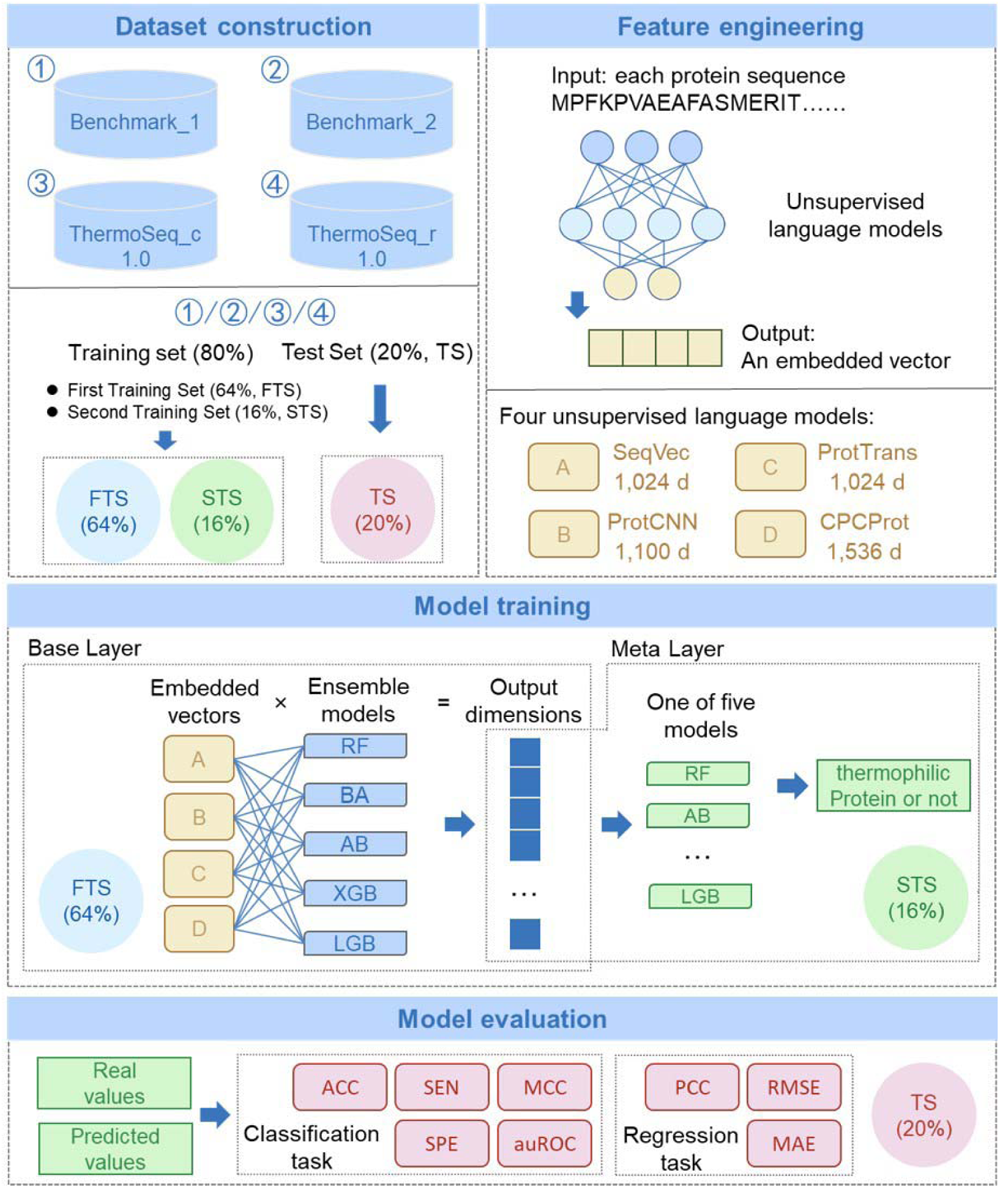
Overview of the ThermoFinder framework. There are four steps: (I) Dataset construction step, (II) Feature engineering step, (III) Model training step, and (IV) Model evaluation step.

#### 2.3.1 Dataset construction step

Each of the four datasets was divided into an 80% training set and a 20% independent test set (ITS) as described in previous references [24–25]. To facilitate the training across various layers of the model, the training set is further split into two subsets. The first training set (FTS) constitutes 80% of the original training set (or 64% of the entire dataset) and is used to train the base layer of the model, while the second training set (STS) comprises the remaining 20% of the training set (or 16% of the entire dataset) and is utilized to train the meta layer of the model. The ITS is exclusively employed to independently evaluate the model’s performance.

#### 2.3.2 Feature engineering step

The feature engineering step extracts crucial information from protein sequences for the following machine learning process. We selected four unsupervised language models, including SeqVec, ProtCNN, ProtTrans, and CPCProt, to generate embedded vectors from each protein sequence. These models are sourced from different model architectures, ensuring that different levels of information are extracted. The resulting embedded vectors have dimensions of 1,024, 1,100, 1,024, and 1,536, respectively.

#### 2.3.3 Model training step

The model training step is comprised of a base layer and a meta layer. For the base layer, we selected five representative ensemble models, including Random Forest (RF), bagging (BA), AdaBoost (AB), XGBoost (XGB), and LightGBM (LGB), due to their powerful fitting ability and mechanisms to prevent over-fitting. Each of the four different protein embedded representations were individually applied to each of the five ensemble models, resulting in a set of 20 independent baseline models. Using the FTS, each baseline model is trained individually to obtain a probability value for predicting thermophilic proteins. Overall, the base layer, containing 20 baseline models, generated a 20-dimensional probability vector for every sequence. For the training of meta layer, we utilized the base layer pre-trained with FTS to generate a 20-dimensional probability vector for each protein sequence in the STS, which, together with their corresponding label of ground truth, were then used as the input dataset, Then a representative ensemble model, randomly selected from the aforementioned five models, were used to predict whether the protein is thermophilic or not. In this study, we implemented all ensemble models using sklearn v. 1.1.1, utilizing default parameters. For LightGBM, we employed LGBMClassifier() with gradient-based learning and 100 boosting iterations. XGBoost involved XGBClassifier() with a gradient boosting algorithm and 100 boosting rounds. AdaBoost used AdaBoostClassifier() with the default decision tree base estimator and 50 estimators for boosting. RandomForest employed RandomForestClassifier() with 100 decision trees and “gini” as the split quality criterion. Finally, Bagging used BaggingClassifier() with 10 base estimators employing the decision tree classifier. Given that the model parameters in the sklearn library are already optimized and usually demonstrate good performance, we did not conduct further optimization.

#### 2.3.4 Model evaluation step

To assess the performance of the model, we employed representative evaluation metrics to compare the proposed tool with current SOTA tools on the ITS. For classification task, we chose to evaluate the accuracy (ACC), sensitivity (SEN), specificity (SPE), area under the receiver operating characteristic curve (auROC), and Matthews correlation coefficient (MCC). Of these metrics, auROC and MCC are considered to be the most representative. We calculated these metrics using the following formulas:

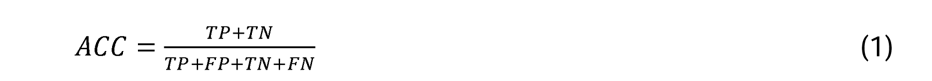

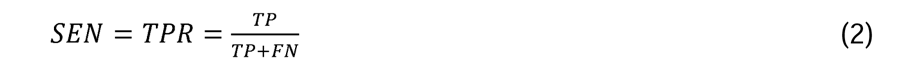

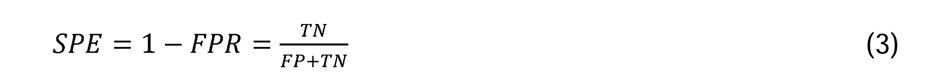

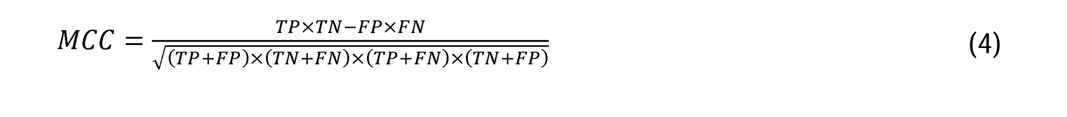

where TP represents the number of correctly predicted positive samples, TN represents the number of correctly predicted negative samples, FP represents the number of negative samples predicted to be positive samples, and FN represents the number of positive samples predicted to be negative samples. The auROC is the area under the curve, which is constructed by considering the FPR as the horizontal axis and the TPR as the vertical axis by setting a different threshold.

For regression task, we calculated pearson correlation coefficient (PCC), root mean square error (RMSE) and mean absolute error (MAE). These metrics were calculated as:

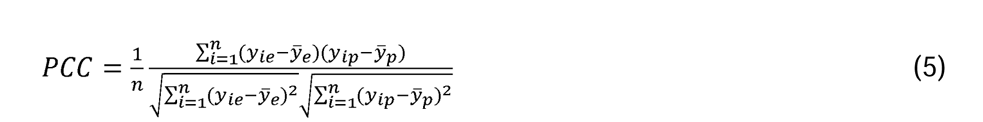

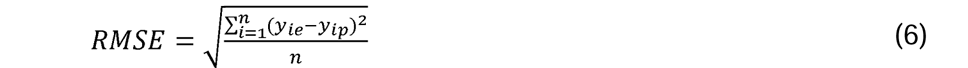

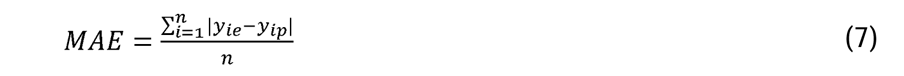

where y_ie_ for the real temperature optimum value, y_ip_ for the predicted temperature optimum value, y _e_ for the average of the real temperature optimum values, y _p_ for the average of the predicted temperature optimum values, and n for the number of samples.

## 3. Results

### 3.1 Comparison between ThermoFinder and existing tools on two benchmark datasets

To assess the effectiveness of our proposed ThermoFinder, we compared its performance to that of existing tools on two benchmark datasets. Specifically, we conducted a performance evaluation of ThermoFinder using five distinct ensemble models in the meta layer. Subsequently, we selected two previously developed representative tools for the same task, iThermo and SCMTPP, for a comparative analysis [24–25]. In addition, we also compared our framework with SAPPHIRE and IPPF-FE, a stacking-based ensemble learning framework for the prediction of thermophilic proteins, and a previously reported state-of-the-art (SOTA) peptide and protein function prediction framework [33].

For the first benchmark dataset, Table 3 shows that all five tools derived from the ThermoFinder framework outperformed iThermo across all metrics in the test set. The XGBoost (XGB) emerged as the optimal choice for the meta layer. The resulting ThermoFinder-XGB achieved higher MCC and auROC values than iThermo by 5.18% and 1.15%, respectively. Moreover, ThermoFinder-XGB exhibited a 2.67% advantage in ACC, with SEN and SPE values of 0.9927 and 0.9862, respectively. The other tools also demonstrated advantages across these metrics in Table 3. Similarly, the results for the second benchmark dataset in Table 4 indicated that all five tools had a significant advantage over various metrics in the test set. ThermoFinder-RF, with random forest (RF) for the meta layer, achieved the best performance, with MCC and auROC values 22.32% and 7.30% higher than SCMTPP, respectively. The superiority of ThermoFinder-RF is further emphasized by additional metrics, affirming its absolute dominance as shown in Table 4. In comparison, while SAPPHIRE, another predictive tool, has displayed accurate predictions on the second benchmark dataset, it hasn’t been able to surpass ThermoFinder-RF’s performance, as evidenced in Table 4.

**Table 3.** Comparison of metric performance among different tools on the independent set for the first benchmark dataset, including ACC, SEN, SPE, MCC, and auROC.

| Tools | Base layer | Meta layer | ACC (%) | SEN (%) | SPE (%) | MCC (%) | auROC (%) |
| --- | --- | --- | --- | --- | --- | --- | --- |
| ThermoFinder-LGB | Five models | LGB | 98.40 | **99.27** | 97.58 | 96.81 | 99.71 |
| ThermoFinder-XGB | Five models | XGB | **98.93** | **99.27** | **98.62** | **97.87** | **99.79** |
| ThermoFinder-AB | Five models | AB | 98.58 | **99.27** | 97.92 | 97.16 | 99.47 |
| ThermoFinder-RF | Five models | RF | 98.58 | **99.27** | 97.92 | 97.16 | 99.72 |
| ThermoFinder-BA | Five models | BA | 98.40 | 98.90 | 97.92 | 96.80 | 99.23 |
| iThermo | - | - | 96.26 | 96.34 | 96.19 | 92.69 | 98.64 |

**Table 4.** Comparison of metric performance among different tools on the independent set for the second benchmark dataset, including ACC, SEN, SPE, MCC, and auROC.

| Tools | Base layer | Meta layer | ACC (%) | SEN (%) | SPE (%) | MCC (%) | auROC (%) |
| --- | --- | --- | --- | --- | --- | --- | --- |
| ThermoFinder-LGB | Five models | LGB | 97.44 | 97.02 | 97.86 | 94.88 | 99.73 |
| ThermoFinder-XGB | Five models | XGB | 97.57 | **97.56** | 97.59 | 95.15 | 99.66 |
| ThermoFinder-AB | Five models | AB | 97.30 | 97.02 | 97.59 | 94.61 | 99.72 |
| ThermoFinder-RF | Five models | RF | **97.71** | **97.56** | 97.86 | **95.42** | **99.80** |
| ThermoFinder-BA | Five models | BA | 97.57 | 97.02 | **98.12** | 95.15 | 98.90 |
| SCMTPP | - | - | 86.50 | 84.90 | 88.10 | 73.10 | 92.50 |
| SAPPHIRE | - | - | 94.20 | 95.10 | 93.30 | 88.40 | 98.00 |

Furthermore, we validated these datasets on IPPF-FE in Supplementary Table S1. Although IPPF-FE demonstrated significant better prediction performance with MCC values of 0.9644 and 0.9383 on these two datasets, respectively, surpassing the performance of iThermo and SCMTPP tools, ThermoFinder-XGB and ThermoFinder-RF still outperformed it. Overall, our results demonstrated that ThermoFinder significantly outperformed previous tools in terms of various metrics, and its comparison with the SOTA protein prediction framework further verified its effectiveness. The next step was to validate the proposed framework’s effectiveness by comparing it with single features.

### 3.2 Comparison of single and fused representations

To confirm the superiority of fused representations, we compared the performance of four single representations with fused representations on the test sets of both benchmark datasets. Here, as mentioned in the previous section, for the first and second benchmark datasets, respectively, the models used were XGB and RF. Our results on the first benchmark dataset showed that ThermoFinder-XGB outperformed single representations across various metrics, including ACC, SEN, SPE, MCC, and auROC. Our model achieved improvements of 1.42-17.44%, 2.56-18.68%, 0.35-16.26%, 2.84-34.91%, and -0.04-9.62% over single representations in Figure 2a. On the second benchmark dataset, ThermoFinder-RF achieved improvements of 1.07-20.35%, 3.49-21.82%, -1.34-18.88%, 2.03-40.67%, and 0.18-16.95% over single representations in Figure 2b. Significantly, the ProtTrans representation emerged as the most crucial and dominant feature. These results confirmed that the remarkable advantages of fused representations introduced by ThermoFinder over single ones. As a next step, to further demonstrate the indispensability of fused representations, we conducted ablation studies.

**Figure 2.**
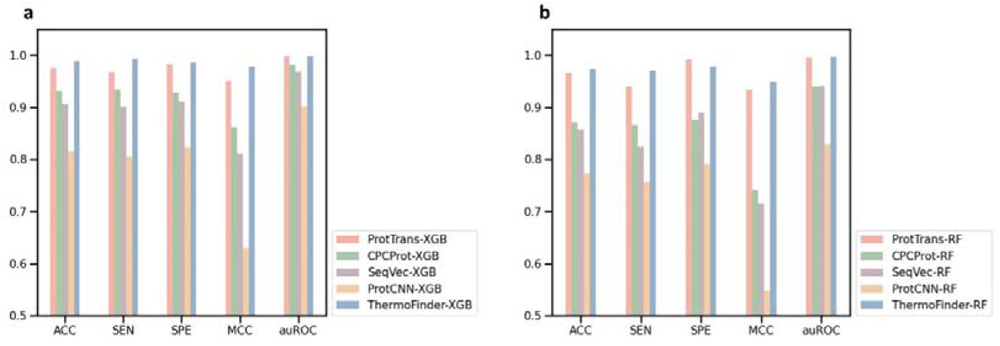
a, b) The metrics comparison of single and fused representations on the independent set in terms of ACC, SEN, SPE MCC and auROC for the first benchmark dataset (a), and second benchmark dataset (b), respectively.

### 3.3 Ablation studies

To further demonstrate the advantages of the proposed ThermoFinder, we conducted a comprehensive ablation study to the base layer. Specifically, we ensembled each of the four individual protein sequence representation with all models, resulted in four different ensembles (I-R) as the base layer. In addition, we ensembled each of the five individual models with all representations, resulted in five different ensembles (I-M) as the base layer, as indicated in Figure 3. The meta layer was identical to ThermoFinder, from which one of the five individual models will be used for the final prediction. Therefore, we obtained a total of 20 (4 I-Rs as base layer * 5 models as meta layer) and 25 (5 I-Ms as base layer * 5 models as meta layer) combinations, respectively.

**Figure 3.**
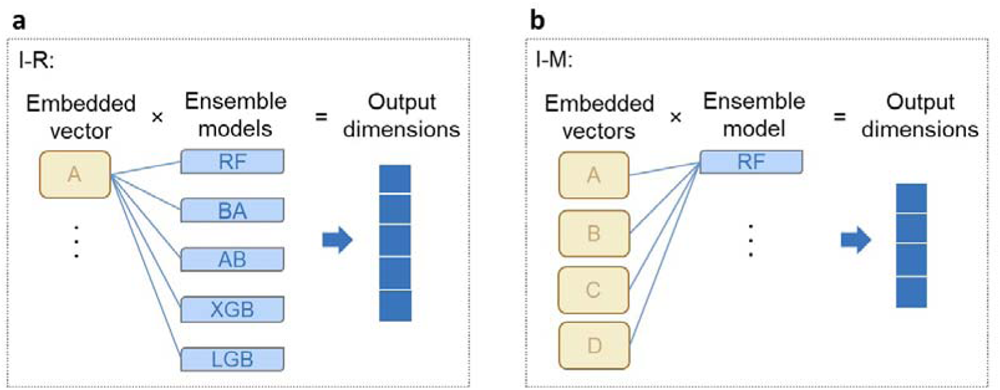
a, b) Overview of the individual protein sequence representation with all models (I-R) (a) and individual models with all representations (I-M) (b) as the base layer.

Regarding I-Rs, our results on the first benchmark dataset showed that the 20 combinations yielded results that were worse than ThermoFinder-XGB, as shown in Supplementary Table S2. The best result was achieved using ProtTrans representation with five models on the base layer and bagging (BA) on the meta layer, with ACC, SEN, SPE, MCC, and auROC values of 0.9858, 0.9817, 0.9896, 0.9715, and 0.9975, respectively. On the second benchmark dataset, ThermoFinder-RF outperformed all 20 combinations in Supplementary Table S3. Again, the best result was achieved using ProtTrans representation with five models on the base layer and RF on the meta layer, with ACC, SEN, SPE, MCC, and auROC values of 0.9717, 0.9675, 0.9759, 0.9434, and 0.9966, respectively.

Regarding I-Ms, the results showed that the prediction performance of all 25 combinations on the first or second benchmark dataset were worse to ThermoFinder-XGB or ThermoFinder-RF, respectively, in Supplementary Table S4-5. The best result was achieved using LightGBM (LGB) on the base layer and RF on the meta layer for the first benchmark dataset, and LGB on both the base and meta layers for the second benchmark dataset. These results provide further evidence of the effectiveness of ThermoFinder, with ProtTrans representation contributing the most to the model’s success.

### 3.4 Performance of ThermoFinder on two newly constructed datasets

To overcome the limitation of current datasets and promote the development of downstream applications, we established two new datasets for classification and regression tasks, named ThermoSeq_c 1.0 and ThermoSeq_r 1.0, respectively. These datasets comprise 63,880 and 7,348 samples, respectively, significantly larger than the previous dataset.

For the ThermoSeq_c 1.0 dataset, we randomly divided the dataset into an 80% training set and a 20% test set. ThermoFinder-RF demonstrated excellent performance on this dataset, with an MCC of 0.8974 and an auROC of 0.9894. We also evaluated two state-of-the-art tools, iThermo and SCMTPP [24–25], on this dataset and found that all ThermoFinder models outperformed them with significantly higher ACC, SEN, SPE, MCC, and auROC values, as presented in Table 5. These results confirmed the effectiveness of ThermoFinder in classification task. For the ThermoSeq_r 1.0 dataset, the division was identical to that of ThermoSeq_c 1.0. The results, as presented in Table 6, revealed that the ThermoFinder-RF emerged as the optimal tool, showcasing a root mean squared error (RMSE) of 11.46 and a mean absolute error (MAE) of 8.42. Furthermore, a notable correlation of 0.70 was observed between the predicted and experimentally measured temperature optimum values.

**Table 5.** Comparison of metric performance among different tools on the independent set for the ThermoSeq_c 1.0 dataset, including ACC, SEN, SPE, MCC, and auROC.

| Tools | Base layer | Meta layer | ACC (%) | SEN (%) | SPE (%) | MCC (%) | auROC (%) |
| --- | --- | --- | --- | --- | --- | --- | --- |
| ThermoFinder-LGB | Five models | LGB | 94.63 | 94.27 | 95.00 | 89.26 | **98.98** |
| ThermoFinder-XGB | Five models | XGB | 94.47 | 94.20 | 94.74 | 88.95 | 98.90 |
| ThermoFinder-AB | Five models | AB | 94.72 | **94.59** | 94.84 | 89.43 | 98.94 |
| ThermoFinder-RF | Five models | RF | **94.87** | 94.30 | **95.44** | **89.74** | 98.94 |
| ThermoFinder-BA | Five models | BA | 94.38 | 93.35 | 95.43 | 88.78 | 98.19 |
| iThermo | - | - | 79.16 | 71.12 | 87.21 | 59.10 | NA |
| SCMTPP | - | - | 71.29 | 52.33 | 90.25 | 46.02 | NA |

Note: NA represents the result is not available.

**Table 6.** Comparison of metric performance among different tools on the independent set for the ThermoSeq_r 1.0 dataset, including RMSE, MAE, and PCC.

| Tools | Base layer | Meta layer | RMSE | MAE | PCC |
| --- | --- | --- | --- | --- | --- |
| ThermoFinder-LGB | Five models | LGB | 11.99 | 8.82 | 0.67 |
| ThermoFinder-XGB | Five models | XGB | 12.23 | 9.05 | 0.66 |
| ThermoFinder-AB | Five models | AB | 12.03 | 8.99 | 0.68 |
| ThermoFinder-RF | Five models | RF | **11.46** | **8.42** | **0.70** |
| ThermoFinder-BA | Five models | BA | 12.01 | 8.88 | 0.67 |

Given that current predictors do not directly predict temperature optimum values, we utilized the two newly constructed datasets to perform an external validation. Specifically, we predicted the probabilities of being thermophilic for all samples in ThermoSeq_r 1.0 using the predictor trained by ThermoSeq_c 1.0 and ranked the data points in descending order according to the predicted probability value. We calculated the average temperature optimum values of the top or the bottom 500, 1000, 2000, 3000, and 3674 samples. The results, as presented in Figure 4, showed that the former had significantly higher temperature optimum values than the latter, confirming a high consistency between the two tools. Overall, these results demonstrate the great performance of ThermoFinder on the two newly constructed datasets and high prediction consistency, highlighting the potential of ThermoFinder in downstream applications.

**Figure 4.**
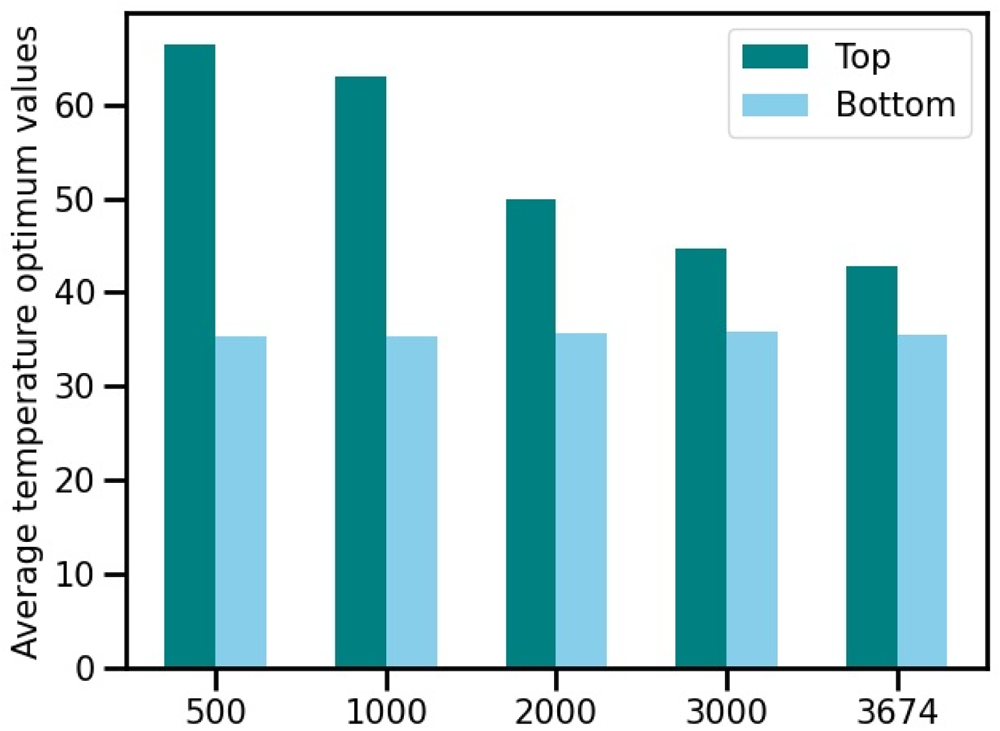
The average temperature optimum values of the top or the bottom 500, 1000, 2000, 3000, and 3674 samples of ThermoSeq_r 1.0 dataset. The predicted probabilities of being thermophilic for all samples in ThermoSeq_r 1.0 using the predictor trained by ThermoSeq_c 1.0 and ranked the data points in descending order according to the predicted probability value.

### 3.5 Model Interpretations

To gain insight into the learning process, we used SHapley Additive exPlanations (SHAP) to analyze feature importance based on ThermoFinder [34]. We evaluated the output of the base layer (the input of the meta layer) for four datasets used in this study: three for classification tasks and one for a regression task. For the first benchmark dataset, we observed that the top five features had dominant effects, including four protTrans representations from different models and one CPCProt representation with XGB, as shown in Figure 5a. The second benchmark dataset showed that the top two features had crucial effects, which were protTrans representations with RF or LGB, as shown in Figure 5b. In the third dataset, ThermoSeq_c 1.0, the dominant feature was also proTtrans with XGB or LGB, as shown in Figure 5c. For the regression task, ThermoSeq_r 1.0, the top four features had dominant effects, including three protTrans representations with AB, RF, or LGB and one SeqVec representation with LGB, as shown in Figure 5d. The protTrans representation plays a vital role in identifying and predicting thermophilic proteins. In addition, XGB and LGB models significantly rank higher among all models and have a more significant impact on the final prediction results. In summary, various datasets rely on diverse protein sequence representations generated by different models, emphasizing the importance for the proposed framework.

**Figure 5.**
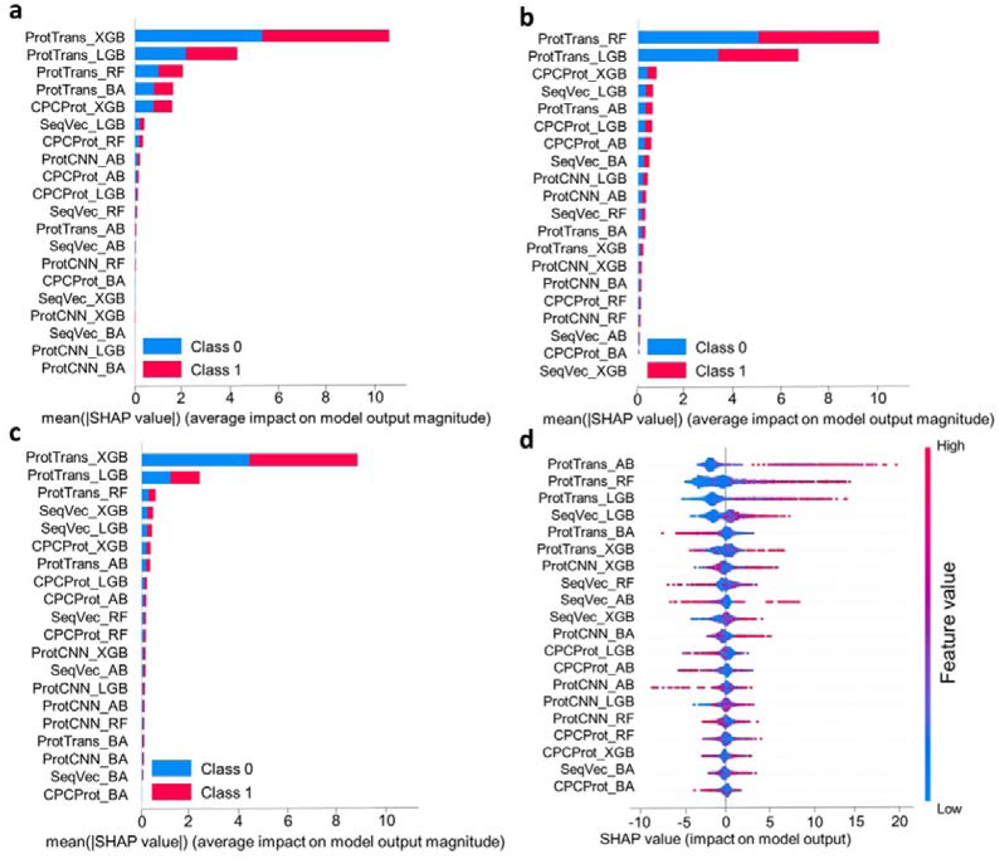
a, b, c, d) Shapley additive explanations (SHAP) analysis for 20 features on the first benchmark dataset (a), second benchmark dataset (b), thermoSeq_c 1.0 dataset (c) and thermoSeq_r 1.0 dataset (d).

## 4 Conclusion

### The conventional tools for predicting thermophilic proteins suffer from

limitations in their datasets and models, which hinder downstream development and application. To address this issue, we proposed a novel sequence-based prediction framework of thermophilic proteins, ThermoFinder, utilizing machine learning technique. We evaluated the effectiveness of our framework on two benchmark datasets, iThermo and SCMTPP, and compared it with existing tools. Our results demonstrated that ThermoFinder surpasses other tools across various metrics, including MCC and auROC. Furthermore, our comparison between single and fused representations confirms the necessity of fused representations in the proposed framework, with the ProtTrans representation being the most crucial and dominant feature among protein sequence representations. Ablation studies further support the effectiveness of ThermoFinder. We speculate that the utilization of multiple pre-trained language models provides a comprehensive approach to represent enzyme sequences, harnessing the strengths of each model and enhancing the overall understanding and analysis of enzyme data. This diverse representation could lead to improved performance in the identification of thermophilic proteins. We also tested our framework on two newly constructed datasets, ThermoSeq_c 1.0 and ThermoSeq_r 1.0, and observed its great performance. To the best of our knowledge, utilizing the ThermoSeq_r 1.0 dataset, we have accomplished the first prediction of temperature optimum values directly from protein sequences, without any additional information. Moreover, we employed SHAP to analyze the importance of features on four datasets. Remarkably, we observed that the protTrans representation significantly influences the identification of thermophilic proteins. Additionally, the XGB and LGB models outperformed other models in our experiments.

In summary, the ThermoFinder framework demonstrated high accuracy and scalability in predicting thermophilic proteins, making it a promising tool for enzyme evolution and engineering. This study is the most comprehensive research to date on predicting thermophilic proteins. However, the model’s predictive power is limited by its reliance on training data, which may not accurately predict unknown types of thermophilic proteins. To improve the model’s generalization capabilities, future research could focus on the information of atom level and capture more general rules [35–36]. Furthermore, we used four unsupervised language models to generate protein sequence representations, which are incomplete given the availability of new representations like variational autoencoder and restricted Boltzmann machine [37–40]. Therefore, we intend to explore these new representations to enhance the identification of thermophilic proteins and their downstream applications in enzyme design and engineering.

## Acknowledgements

This work was supported by the National Key R&D Program of China (2018YFA0903200), National Natural Science Foundation of China (32071421), Guangdong Basic and Applied Basic Research Foundation (2021B1515020049), Shenzhen Science and Technology Program (ZDSYS20210623091810032 and JCYJ20220531100207017), and Shenzhen Institute of Synthetic Biology Scientific Research Program (ZTXM20203001). We also want to thank Miss Z. Wei for the support in handling administrative affairs.

## Author contributions

H.Y. and X.L. conceived and designed the study, analyzed and interpreted the data, drafted and revised the manuscript. All authors read and approved the final manuscript.

## Data Availability Statement

The data and model are available on Github at https://github.com/Luo-SynBioLab/ThermoFinder.

## References

[1] Szilágyi A, Závodszky P. Structural differences between mesophilic, moderately thermophilic and extremely thermophilic protein subunits: results of a comprehensive survey. Structure. 2000;8(5):493–504.

[2] Finch AJ, Kim JR. Thermophilic Proteins as Versatile Scaffolds for Protein Engineering. Microorganisms. 2018;6(4):97

[3] Cowan, D, Roy D, and Hugh M. Thermophilic proteases: properties and potential applications. Trends Biotechnol. 1985;3(3):68–72

[4] Blumer-Schuette SE, Brown SD, Sander KB, et al. Thermophilic lignocellulose deconstruction. FEMS Microbiol Rev. 2014;38(3):393–448.

[5] Kumwenda B, Litthauer D, Bishop OT, et al. Analysis of protein thermostability enhancing factors in industrially important thermus bacteria species. Evol Bioinform Online. 2013;9:327–42.

[6] Tang W, Dai R, Yan W, et al. Identifying multi-functional bioactive peptide functions using multi-label deep learning. Brief Bioinform. 2022;23(1):bbab414.

[7] Zhang Y, Lin J, Zhao L, et al. A novel antibacterial peptide recognition algorithm based on BERT. Brief Bioinform. 2021;22(6):bbab200.

[8] Charoenkwan P, Chiangjong W, Nantasenamat C, et al. A large-scale evaluation of computational protein function prediction. Nat Methods. 2013;10(3):221–7.

[9] Kulmanov M, Hoehndorf R. DeepGOPlus: improved protein function prediction from sequence. Bioinformatics. 2021;37(8):1187.

[10] Feng C, Ma Z, Yang D, et al. A Method for Prediction of Thermophilic Protein Based on Reduced Amino Acids and Mixed Features. Front Bioeng Biotechnol. 2020;8:285.

[11] Guo Z, Wang P, Liu Z, et al. Discrimination of Thermophilic Proteins and Non-thermophilic Proteins Using Feature Dimension Reduction. Front Bioeng Biotechnol. 2020;8:584807.

[12] Charoenkwan P, Schaduangrat N, Moni MA, et al. SAPPHIRE: A stacking-based ensemble learning framework for accurate prediction of thermophilic proteins. Comput Biol Med. 2022;146:105704.

[13] Meng C, Ju Y, Shi H. TMPpred: A support vector machine-based thermophilic protein identifier. Anal Biochem. 2022;645:114625.

[14] Zhao J, Yan W, Yang Y. DeepTP: A Deep Learning Model for Thermophilic Protein Prediction. Int J Mol Sci. 2023;24(3):2217.

[15] Liu X, Zhang F, Hou Z, et al. Self-supervised learning: Generative or contrastive. IEEE T Knowl Data En. 2021;35(1), 857–76.

[16] Jaiswal A, Babu AR, Zadeh MZ, et al. A survey on contrastive self-supervised learning. Technologies. 2020;9(1): 2.

[17] Zhai X, Oliver A, Kolesnikov A, et al. S4l: Self-supervised semi-supervised learning. Proceedings of the IEEE/CVF international conference on computer vision 2019;1476–85.

[18] Unsal S, Atas H, Albayrak M, et al. Learning functional properties of proteins with language models. Nat Mach Intell. 2022;4(3), 227-45.

[19] Bileschi ML, Belanger D, Bryant DH, et al. Using deep learning to annotate the protein universe. Nat Biotechnol. 2022;40(6):932–937.

[20] Elnaggar A, Heinzinger M, Dallago C, et al. ProtTrans: Toward Understanding the Language of Life Through Self-Supervised Learning. IEEE Trans Pattern Anal Mach Intell. 2022;44(10):7112–7127.

[21] Teufel F, Almagro AJ, Johansen AR, et al. SignalP 6.0 predicts all five types of signal peptides using protein language models. Nat Biotechnol. 2022;40(7):1023–1025.

[22] Manfredi M, Savojardo C, Martelli PL, et al. E-SNPs&GO: Embedding of protein sequence and function improves the annotation of pathogenic variants. Bioinformatics 2022;btac678.

[23] Singh J, Paliwal K, Singh J, et al. Reaching alignment-profilebased accuracy in predicting protein secondary and tertiary structural properties without alignment. Sci Rep 2022;12(1):1–9.

[24] Ahmed Z, Zulfiqar H, Khan AA, et al. iThermo: A Sequence-Based Model for Identifying Thermophilic Proteins Using a Multi-Feature Fusion Strategy. Front Microbiol. 2022;13:790063.

[25] Charoenkwan P, Chotpatiwetchkul W, Lee VS, et al. A novel sequence-based predictor for identifying and characterizing thermophilic proteins using estimated propensity scores of dipeptides. Sci Rep. 2021;11(1):23782.

[26] Engqvist MKM. Correlating enzyme annotations with a large set of microbial growth temperatures reveals metabolic adaptations to growth at diverse temperatures. BMC Microbiol. 2018;18(1):177.

[27] UniProt Consortium. UniProt: a worldwide hub of protein knowledge. Nucleic Acids Res. 2019;47(D1):D506–15.

[28] Fu L, Niu B, Zhu Z, et al. CD-HIT: accelerated for clustering the next-generation sequencing data. Bioinformatics 2012;28:3150–2.

[29] Schomburg I, Jeske L, Ulbrich M, et al. The BRENDA enzyme information system-From a database to an expert system. J Biotechnol. 2017;261:194–206.

[30] Almagro Armenteros JJ, Sønderby CK, Sønderby SK, et al. DeepLoc: prediction of protein subcellular localization using deep learning. Bioinformatics 2017;33(21):3387–95.

[31] Heinzinger M, Elnaggar A, Wang Y, et al. Modeling aspects of the language of life through transfer-learning protein sequences. BMC Bioinformatics. 2019;20(1):723.

[32] Lu AX, Zhang H, Ghassemi M, et al. Self-supervised contrastive learning of protein representations by mutual information maximization. BioRxiv. 2020.

[33] Yu H, Luo X. IPPF-FE: an integrated peptide and protein function prediction framework based on fused features and ensemble models. Brief Bioinform. 2023;24(1):bbac476.

[34] Lundberg SM, Lee SI. A unified approach to interpreting model predictions. Advances in neural information processing systems, 2017.

[35] King JE, Koes DR. SidechainNet: An all-atom protein structure dataset for machine learning. Proteins: Structure, Function and Bioinformatics. 2021;89(11), 1489–96.

[36] Li B, Yang YT, Capra JA, et al. Predicting changes in protein thermodynamic stability upon point mutation with deep 3D convolutional neural networks. PLoS comput biol. 2020;16(11):e1008291.

[37] Riesselman AJ, Ingraham JB, & Marks DS. Deep generative models of genetic variation capture the effects of mutations. Nat. Methods 2018;15:816–22.

[38] Tubiana J, Cocco S & Monasson R. Learning protein constitutive motifs from sequence data. eLife. 2019;8:e39397.

[39] Alley EC, Khimulya G, Biswas S, et al. Unified rational protein engineering with sequence-based deep representation learning. Nat. Methods. 2019;16:1315–22.

[40] Bepler T, Berger B. Learning protein sequence embeddings using information from structure. 2019.

