## Supplementary Table S1 for "ThermoFinder: A sequence-based thermophilic proteins prediction framework"

Table S1 Comparison of metric performance between ThermoFinder and IPPF-FE on the independent set for two benchmark datasets, including ACC, SEN, SPE, MCC, and auROC.

| Datasets | Tools | ACC (%) | SEN (%) | SPE (%) | MCC (%) | auROC (%) |
| --- | --- | --- | --- | --- | --- | --- |
| Benckmark_1 | ThermoFinder-XGB | **98.93** | **99.27** | **98.62** | **97.87** | 99.79 |
|  | IPPF-FE | 98.22 | 98.17 | 98.27 | 96.44 | **99.84** |
| Benckmark_2 | ThermoFinder-RF | **97.71** | **97.56** | **97.86** | **95.42** | **99.80** |
|  | IPPF-FE | 96.90 | 95.69 | 98.11 | 93.83 | 99.63 |

Table S2 Comparison of metric performance among 20 (4 I-Rs as base layer * 5 models as meta layer) tools on the independent set for the first benchmark dataset, including ACC, SEN, SPE, MCC, and auROC.

| Tools | Base layer | Meta layer | ACC (%) | SEN (%) | SPE (%) | MCC (%) | auROC (%) |
| --- | --- | --- | --- | --- | --- | --- | --- |
| ProtTrans+LGB | Five models | LGB | 98.04 | 97.80 | 98.27 | 96.08 | 99.88 |
| ProtTrans+XGB | Five models | XGB | 98.40 | 97.80 | 98.96 | 96.80 | 99.69 |
| ProtTrans+AB | Five models | AB | 97.86 | 97.44 | 98.27 | 95.73 | 98.94 |
| ProtTrans+RF | Five models | RF | 98.58 | 98.53 | 98.62 | 97.15 | 99.74 |
| ProtTrans+BA | Five models | BA | **98.58** | **98.17** | **98.96** | **97.15** | **99.75** |
| CPCProt+LGB | Five models | LGB | 91.46 | 92.67 | 90.31 | 82.95 | 96.30 |
| CPCProt+XGB | Five models | XGB | 90.75 | 91.94 | 89.62 | 81.53 | 94.92 |
| CPCProt+AB | Five models | AB | 89.15 | 89.38 | 88.93 | 78.29 | 96.17 |
| CPCProt+RF | Five models | RF | 91.99 | 94.87 | 89.27 | 84.15 | 96.62 |
| CPCProt+BA | Five models | BA | 90.75 | 94.14 | 87.54 | 81.72 | 95.79 |
| SeqVec+LGB | Five models | LGB | 89.15 | 89.38 | 88.93 | 78.29 | 94.81 |
| SeqVec+XGB | Five models | XGB | 89.86 | 90.84 | 88.93 | 79.74 | 94.59 |
| SeqVec+AB | Five models | AB | 89.86 | 89.74 | 89.97 | 79.70 | 95.02 |
| SeqVec +RF | Five models | RF | 90.75 | 92.31 | 89.27 | 81.55 | 95.15 |
| SeqVec +BA | Five models | BA | 90.39 | 91.58 | 89.27 | 80.82 | 93.82 |
| ProtCNN+LGB | Five models | LGB | 81.14 | 83.88 | 78.55 | 62.44 | 89.45 |
| ProtCNN+XGB | Five models | XGB | 79.54 | 82.42 | 76.82 | 59.26 | 88.23 |
| ProtCNN+AB | Five models | AB | 81.85 | 83.15 | 80.62 | 63.75 | 89.43 |
| ProtCNN+RF | Five models | RF | 84.88 | 87.18 | 82.70 | 69.87 | 90.34 |
| ProtCNN+BA | Five models | BA | 84.34 | 85.71 | 83.04 | 68.73 | 88.53 |

Table S3 Comparison of metric performance among 20 (4 I-Rs as base layer * 5 models as meta layer) tools on the independent set for the second benchmark dataset, including ACC, SEN, SPE, MCC, and auROC.

| Tools | Base layer | Meta layer | ACC (%) | SEN (%) | SPE (%) | MCC (%) | auROC (%) |
| --- | --- | --- | --- | --- | --- | --- | --- |
| ProtTrans+LGB | Five models | LGB | 96.50 | 96.75 | 96.25 | 92.99 | 99.47 |
| ProtTrans+XGB | Five models | XGB | 97.04 | 97.02 | 97.05 | 94.07 | 99.61 |
| ProtTrans+AB | Five models | AB | 97.04 | 96.75 | 97.32 | 94.07 | 99.51 |
| ProtTrans+RF | Five models | RF | **97.17** | **96.75** | **97.59** | **94.34** | **99.66** |
| ProtTrans+BA | Five models | BA | 97.17 | 97.02 | 97.32 | 94.34 | 99.14 |
| CPCProt+LGB | Five models | LGB | 86.66 | 88.62 | 84.72 | 73.38 | 94.16 |
| CPCProt+XGB | Five models | XGB | 85.44 | 87.26 | 83.65 | 70.95 | 93.36 |
| CPCProt+AB | Five models | AB | 84.23 | 89.16 | 79.36 | 68.82 | 93.82 |
| CPCProt+RF | Five models | RF | 86.52 | 90.24 | 82.84 | 73.26 | 94.65 |
| CPCProt+BA | Five models | BA | 82.75 | 82.11 | 83.38 | 65.50 | 92.42 |
| SeqVec+LGB | Five models | LGB | 86.66 | 86.99 | 86.33 | 73.32 | 94.49 |
| SeqVec+XGB | Five models | XGB | 86.39 | 86.45 | 86.33 | 72.78 | 93.75 |
| SeqVec+AB | Five models | AB | 86.12 | 86.45 | 85.79 | 72.24 | 94.34 |
| SeqVec +RF | Five models | RF | 87.87 | 87.26 | 88.47 | 75.74 | 95.21 |
| SeqVec +BA | Five models | BA | 87.33 | 86.72 | 87.94 | 74.67 | 94.40 |
| ProtCNN+LGB | Five models | LGB | 75.07 | 79.67 | 70.51 | 50.38 | 82.01 |
| ProtCNN+XGB | Five models | XGB | 73.72 | 76.69 | 70.78 | 47.55 | 80.21 |
| ProtCNN+AB | Five models | AB | 74.53 | 79.67 | 69.44 | 49.36 | 83.61 |
| ProtCNN+RF | Five models | RF | 76.95 | 80.49 | 73.46 | 54.07 | 83.63 |
| ProtCNN+BA | Five models | BA | 74.26 | 72.63 | 75.87 | 48.53 | 80.26 |

Table S4 Comparison of metric performance among 25 (5 I-Ms as base layer * 5 models as meta layer) tools on the independent set for the first benchmark dataset, including ACC, SEN, SPE, MCC, and auROC.

| Tools | Base layer | Meta layer | ACC (%) | SEN (%) | SPE (%) | MCC (%) | auROC (%) |
| --- | --- | --- | --- | --- | --- | --- | --- |
| LGB+LGB | LGB | LGB | 98.40 | 98.90 | 97.92 | 96.80 | 99.79 |
| LGB+XGB | LGB | XGB | 98.40 | 98.90 | 97.92 | 96.80 | 99.73 |
| LGB+AB | LGB | AB | 98.40 | 99.27 | 97.58 | 96.81 | 99.67 |
| LGB+RF | LGB | RF | **98.58** | **98.90** | **98.27** | **97.15** | **99.44** |
| LGB+BA | LGB | BA | 98.22 | 98.90 | 97.58 | 96.45 | 99.38 |
| XGB+LGB | XGB | LGB | 98.22 | 98.90 | 97.58 | 96.45 | 99.28 |
| XGB+XGB | XGB | XGB | 98.40 | 98.90 | 97.92 | 96.80 | 99.51 |
| XGB+AB | XGB | AB | 98.40 | 98.90 | 97.92 | 96.80 | 99.49 |
| XGB+RF | XGB | RF | 98.40 | 98.90 | 97.92 | 96.80 | 99.42 |
| XGB+BA | XGB | BA | 98.40 | 98.17 | 98.62 | 96.79 | 99.23 |
| AB+LGB | AB | LGB | 98.04 | 98.17 | 97.92 | 96.08 | 99.85 |
| AB+XGB | AB | XGB | 98.04 | 98.17 | 97.92 | 96.08 | 99.85 |
| AB+AB | AB | AB | 97.33 | 97.44 | 97.23 | 94.66 | 99.65 |
| AB+RF | AB | RF | 98.22 | 98.90 | 97.58 | 96.45 | 99.90 |
| AB+BA | AB | BA | 98.58 | 98.90 | 98.27 | 97.15 | 99.58 |
| RF+LGB | RF | LGB | 97.86 | 97.80 | 97.92 | 95.73 | 99.23 |
| RF+XGB | RF | XGB | 98.04 | 97.44 | 98.62 | 96.09 | 99.18 |
| RF+AB | RF | AB | 97.15 | 97.80 | 96.54 | 94.31 | 99.42 |
| RF+RF | RF | RF | 98.04 | 97.07 | 98.96 | 96.10 | 99.15 |
| RF+BA | RF | BA | 97.86 | 97.44 | 98.27 | 95.73 | 98.65 |
| BA+LGB | BA | LGB | 95.73 | 95.60 | 95.85 | 91.45 | 98.86 |
| BA+XGB | BA | XGB | 95.37 | 95.97 | 94.81 | 90.75 | 98.96 |
| BA+AB | BA | AB | 95.55 | 95.60 | 95.50 | 91.10 | 98.64 |
| BA+RF | BA | RF | 95.73 | 95.60 | 95.85 | 91.45 | 99.14 |
| BA+BA | BA | BA | 95.55 | 94.87 | 96.19 | 91.10 | 98.37 |

Table S5 Comparison of metric performance among 25 (5 I-Ms as base layer * 5 models as meta layer) tools on the independent set for the second benchmark dataset, including ACC, SEN, SPE, MCC, and auROC.

| Tools | Base layer | Meta layer | ACC (%) | SEN (%) | SPE (%) | MCC (%) | auROC (%) |
| --- | --- | --- | --- | --- | --- | --- | --- |
| LGB+LGB | LGB | LGB | **97.84** | **97.29** | **98.39** | **95.69** | **99.80** |
| LGB+XGB | LGB | XGB | 97.44 | 97.29 | 97.59 | 94.88 | 99.75 |
| LGB+AB | LGB | AB | 97.17 | 97.29 | 97.05 | 94.34 | 99.77 |
| LGB+RF | LGB | RF | 97.44 | 97.56 | 97.32 | 94.88 | 99.61 |
| LGB+BA | LGB | BA | 97.04 | 97.02 | 97.05 | 94.07 | 98.89 |
| XGB+LGB | XGB | LGB | 96.63 | 96.21 | 97.05 | 93.26 | 99.75 |
| XGB+XGB | XGB | XGB | 96.77 | 95.93 | 97.59 | 93.54 | 99.69 |
| XGB+AB | XGB | AB | 97.04 | 95.93 | 98.12 | 94.09 | 99.72 |
| XGB+RF | XGB | RF | 96.77 | 96.21 | 97.32 | 93.54 | 99.71 |
| XGB+BA | XGB | BA | 97.04 | 96.48 | 97.59 | 94.08 | 99.00 |
| AB+LGB | AB | LGB | 97.04 | 96.21 | 97.86 | 94.08 | 99.71 |
| AB+XGB | AB | XGB | 97.30 | 95.93 | 98.66 | 94.64 | 99.74 |
| AB+AB | AB | AB | 97.17 | 97.29 | 97.05 | 94.34 | 99.61 |
| AB+RF | AB | RF | 97.04 | 95.93 | 98.12 | 94.09 | 99.66 |
| AB+BA | AB | BA | 97.17 | 96.48 | 97.86 | 94.35 | 99.39 |
| RF+LGB | RF | LGB | 96.63 | 95.93 | 97.32 | 93.27 | 99.42 |
| RF+XGB | RF | XGB | 96.50 | 95.66 | 97.32 | 93.00 | 99.42 |
| RF+AB | RF | AB | 96.50 | 95.12 | 97.86 | 93.02 | 99.64 |
| RF+RF | RF | RF | 96.90 | 95.93 | 97.86 | 93.82 | 99.54 |
| RF+BA | RF | BA | 96.77 | 95.39 | 98.12 | 93.56 | 98.97 |
| BA+LGB | BA | LGB | 95.42 | 94.85 | 95.98 | 90.84 | 99.22 |
| BA+XGB | BA | XGB | 95.82 | 95.39 | 96.25 | 91.65 | 99.02 |
| BA+AB | BA | AB | 95.42 | 95.66 | 95.17 | 90.84 | 99.04 |
| BA+RF | BA | RF | 96.23 | 95.39 | 97.05 | 92.46 | 99.40 |
| BA+BA | BA | BA | 95.55 | 95.39 | 95.71 | 91.11 | 98.43 |
